# Neural differentiation is increased by GSK-3β inhibition and decreased by tankyrase inhibition in human neural precursor cells

**DOI:** 10.1101/509638

**Authors:** Michael Telias, Dalit Ben-Yosef

## Abstract

Glycogen synthase kinase-3β (GSK-3β) and tankyrase-1/2 (TANK) are two enzymes known to play multiple roles in cell biology, including regulation of proliferation, differentiation and metabolism. Both of them act on the canonical Wnt/β-Catenin pathway, but are also involved in many other independent intracellular mechanisms. More importantly, GSK-3β and TANK have been shown to play crucial roles in different diseases, including cancer and neurological disorders. The GSK-3β-inhibitor ‘CHIR’ and the TANK-inhibitor ‘XAV’ are two pyrimidine molecules, holding high potential as possible therapeutic drugs. However, their effect on neural tissue is poorly understood. In this study, we tested the effects of CHIR and XAV on human neural precursor cells (hNPCs) derived from human embryonic stem cells. We found that CHIR-mediated inhibition of GSK-3β promotes neural differentiation. In contrast, XAV-mediated inhibition of TANK leads to de-differentiation. These results highlight the relative importance of these two enzymes in determining the neurodevelopmental status of hNPCs. Furthermore, they shed light on the roles of Wnt signaling during early human neurogenesis.

## INTRODUCTION

The canonical Wnt/ β-Catenin signaling pathway has been shown to play pivotal roles in embryonic neural development as well as in adult neurogenesis [1, 2]. For example, aberrant regulation of Wnt signaling has been proposed to underlie some of the symptoms observed in Alzheimer’s disease, Autism and Fragile X Syndrome [3–8]. Activation of the pathway takes place when the Wnt ligand binds to its receptor, triggering the release of β-Catenin from its inhibitory complex, which enables β-Catenin to translocate to the nucleus, activating or repressing the expression of several target genes [9–11]. Glycogen synthase kinase-3β (GSK-3β) and tankyrase-1/2 (TANK) are two enzymes known to play multiple roles in cell biology, including proliferation, differentiation and metabolism [12–15]. Although each of these enzymes is involved in many independent intracellular mechanisms, they are both connected to the canonical Wnt signaling pathway through regulation of β-Catenin activity, in opposite directions: while GSK-3β activity reduces β-Catenin levels and down-regulates Wnt signaling, TANK activity increases β-Catenin levels and up-regulates Wnt signaling. For these reasons, much effort has been invested in the last years in developing suitable small molecules that could inhibit or modulate the enzymatic activity of GSK-3β and TANK [16–18]. The amino-pyrimidine CHIR99021 (‘CHIR’) is a highly specific inhibitor of GSK-3β, with promising applications as a therapeutic drug [19]. The pyrimidine XAV939 (‘XAV’) is a specific inhibitor of TANK and has also been successfully tested as a promising therapeutic drug [20–22].

Currently, most studies demonstrating the importance of Wnt in neural development have focused mainly on animal-based models and in adult neurogenesis. Reports on the effects of CHIR and XAV so far have also been predominantly performed on animal models, and in non-neural tissues. Recently, we reported on the molecular mechanisms regulating impaired in-vitro neural differentiation of Fragile X Syndrome (FXS)-human embryonic stem cells (hESCs), carrying the full mutation at the FMR1 gene [8]. In that study, we hypothesized that dysregulation of Wnt signaling could be responsible for the poor neural differentiation associated with these cells [23–25]. To test that hypothesis, we applied CHIR or XAV to these cells while carrying out neuronal differentiation. Contrary to what was reported in other FXS models, mainly adult neurogenesis in *fmr1^-/-^* mice hippocampi [4], we observed that there was no significant difference between FX-hESCs and controls. However, and most importantly, we did notice that in all lines tested, CHIR increased neural differentiation while XAV decreased it.

This interesting observation prompted us to further explore the effects of pharmacological manipulation of Wnt on the neural fate of hESCs. Neural differentiation of hESCs is a powerful tool in the study of neurodevelopment in a human-based in-vitro platform [24, 26, 27]. Here, we analyzed the effects of CHIR and XAV on human neural precursor cells (hNPCs) derived from control non-mutated hESC lines. These cells are initially differentiated from hESCs but, under specific culture conditions, can be kept as hNPCs for up to 10 passages [8]. During that time, they can be induced to differentiate into neurons on-cue, by changing their culture conditions, removing basic fibroblast growth factor (bFGF) and adding neuronal growth factors [8, 23, 24]. In this study we induced neuronal differentiation of hNPCs, while adding CHIR or XAV. Our results show that CHIR is a potent neuralizing agent, whereas XAV induces de-differentiation of hNPCs. Importantly, our data are obtained from in-vitro bioassays developed specifically for the measurement of the relative effect of these two inhibitors.

## MATERIALS AND METHODS

### Human embryonic stem cell lines

Four non-affected human embryonic stem cells (hESC) lines were used: HUES-6, HUES-16, HUES-13 and HUES-64 [8, 28-30]. Cells were cultured on Matrigel (BD), in hESC medium supplemented with bFGF (8ng/ml, R&D) [24]. For full information on these cell lines, see Table 1.

### In-vitro Neural Differentiation of hESCs and derivation of hNPCs

In-vitro neural differentiation (IVND) of hESCs was carried out as previously described [8, 24]. In brief, hESCs were grown in Neural Induction Medium (NIM) consisting of DMEM:F12 (LifeTech.), 0.5% B27 (LifeTech.), 1% N2 (LifeTech.), 1% Glutamax (LifeTech.), 1% Non-essential amino acids (BioInd.) and 0.1mg/ml Primocin (InvivoGen). IVND included 3 steps: **(a)** formation of neuro-ectoderm aggregates in the presence of noggin (250ng/ml, PeproTech) and bFGF (20ng/ml, R&E); **(b)** development of attached neural rosettes in the presence of Shh (200ng/ml, PeproTech) and **(c)** formation of neurospheres NIM supplemented with bFGF (20ng/ml, R&E). Human Neural Precursor Cells (hNPCs) were isolated during IVND of hESCs, following mechanical removal of neural rosettes. hNPCs were cultured on Matrigel (BD)-coated polystyrene wells in NIM supplemented with bFGF (20ng/ml). To induce neuronal differentiation, hNPCs were dissociated using TryplE (LifeTech.) and re-plated on Poly-D-Lysine/Laminin (Sigma)-coated glass coverslips. NIM was changed to Neuronal Differentiation Medium (NDM) supplemented with BDNF, GDNF and NT-3 (10ng/ml, PeproTech). NDM consisted of Neurobasal (LifeTech.), 1% B27 (LifeTech.), 1% N2 (Life Tech.), 1% Glutamax (LifeTech.), 1% Non-essential amino acids (BioInd.) and 0.1mg/ml Primocin (InvivoGen).

### Gene transcription analysis

Relative transcription levels were analyzed by quantitative RT-PCR, as previously described [8, 24]. RNA was extracted (RNeasy, Qiagen), reversed transcribed using Super Script-III kit (Invitrogen), and analyzed using SYBRgreen (ABgene) in Rotor Gene 6000 Series (Corbett). The house keeping gene GAPDH was used as a control for ΔΔCt analysis. All qRT-PCR assays included non-template control, non-human cells (MEF), and human-FXS white blood cells. For all primers used see **Table S1**.

### Immunostaining Assay

Immunostaining was performed as previously described [8, 24]. Cells were seeded on glass coverslips coated with Matrigel for hNPCs or with Poly-D-Lysine/Laminin for neuronal differentiation. Cells were fixated using Cytofix (BD). Incubation with primary antibodies was performed over-night at 4°C, detected using Cy2/Cy3-conjugated secondary antibodies. Nuclei were stained using DAPI (Sigma). Coverslips were mounted using Fluoromount G (Southern Biotech). Cells were imaged using an inverted fluorescent microscope (Olympus). All conditions were similar for all lines in all experiments. Each experiment was performed in triplicates and images were taken from 5 different fields/coverslip and >50 cells/field were analyzed by manual counting of positively stained cells and by measurement of mean gray value of Cy2/Cy3 staining relative to DAPI, using ImageJ software (NIH). For all antibodies used see **Table S2**.

### Western Blot Analysis

Western Blot analysis was carried out as previously described [8]. Protein was extracted using reporter lysis buffer (Promega), and 25-30 μg of protein were loaded on a 7.5% separating gel using Mini Trans-Blot Cell (Bio-Rad). Nitrocellulose membranes were stained with primary antibodies, and detected with HRP-conjugated secondary antibodies. Protein bends were detected using EZ-ECL (BioInd.)

### Pharmacological manipulation of Wnt signaling

The GSK3β inhibitor CHIR99021 (Tocris) and the tankyrase inhibitor XAV939 (Selleckchem) were dissolved in DMSO at 20 mM stock solutions and stored at −80°C, in the dark. Fresh CHIR/XAV was added with every medium change, every 48 hrs. Working concentrations of both inhibitors was 3μM in all experiments, and 0.015% DMSO in culture medium.

### Proliferation and Survival Assay

Cells were seeded at low densities (~30,000 cells p/well in a 12-well plate) and incubated for 7 days with either CHIR99021 or XAV939. For assessment of proliferation, cells were immunostained against the proliferation marker Ki-67 (R&D), and quantified relative to DAPI staining. For survival assay, cells were manually counted using a standard hemocytometer

### Statistical Analysis

Statistical analysis (Student’s t-test and ANOVA) was performed using SPSS, SigmaPlot and online GraphPad QuickCalcs (http://www.graphpad.com/quickcalcs).

## RESULTS

### Immediate effects of Wnt modulators on hNPCs

In the present study we have tested the effects of CHIR, a specific GSK-3β inhibitor and XAV, a specific TANK inhibitor, on gene expression, morphology and differentiation potential of hNPCs lines we have previously derived from 4 different hESC lines (Table 1).

**Table 1:** List of all hESC lines used in the present study, its parent institution, its sex and references for further information.

| hESC<br>Line | Institution | Sex | Sources/Information |
| --- | --- | --- | --- |
| HUES6 | Harvard University | XX | http://stemcelldistribution.harvard.edu/<br>[8, 24] |
| HUES16 | Harvard University | XY | http://stemcelldistribution.harvard.edu/<br>[8, 24] |
| HUES13 | Harvard University | XY | http://stemcelldistribution.harvard.edu/<br>[8, 23-25, 27] |
| HUES64 | Harvard University | XY | http://stemcelldistribution.harvard.edu/<br>[8] |

The effects of CHIR and XAV on neuronal differentiation of hNPCs were analyzed following short-term (2-3 days), mid-term (7 days), or long-term treatment (30 days) (Figure 1A).

**Figure 1.**
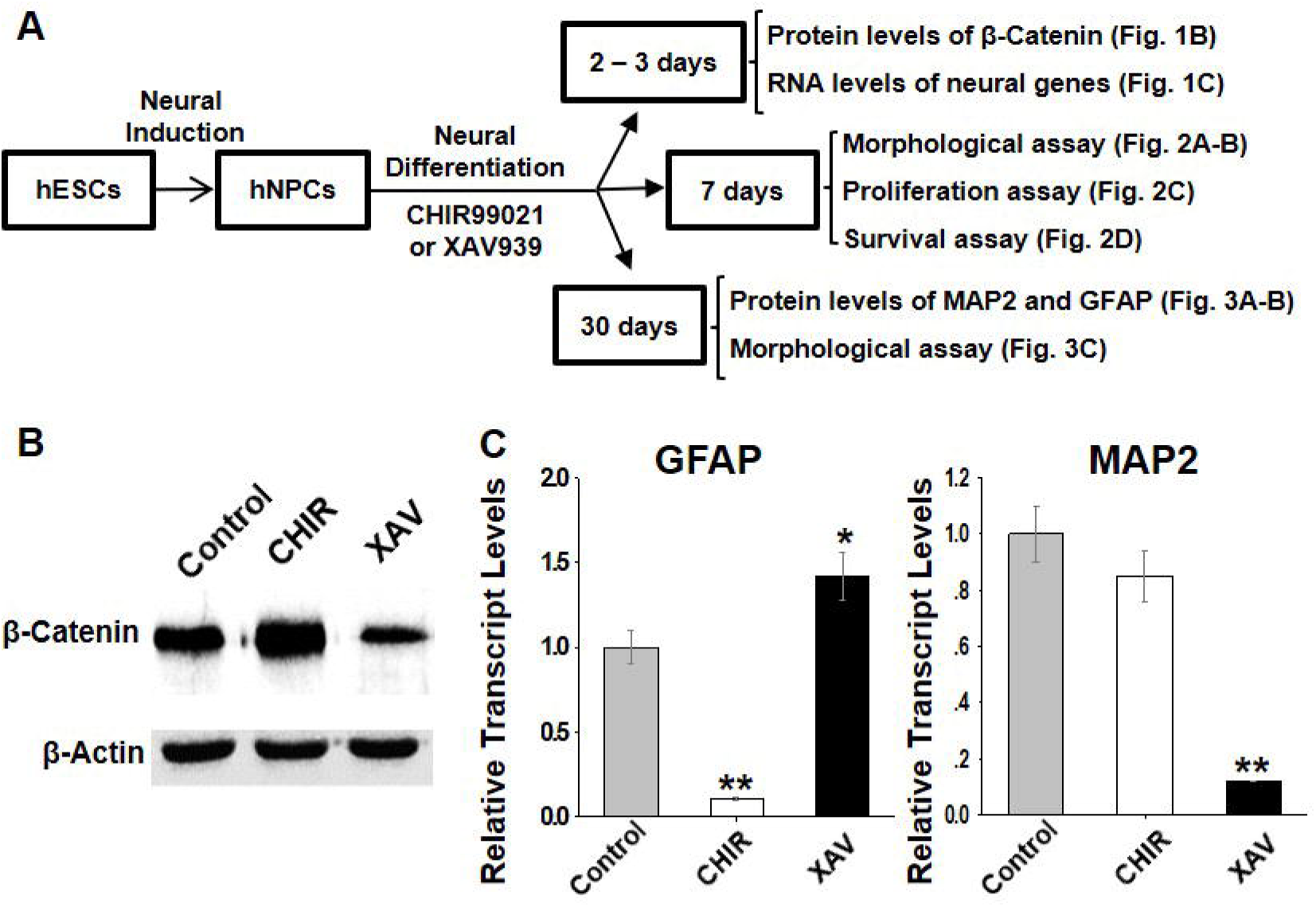
The effect of Wnt modulators on neural gene expression in hNPCs derived from hESCs. **(A)** Schematic outline of the experimental set-up of the present study. **(B)** The effect of CHIR and XAV (3 μM, 2-3 days) on β-Catenin protein levels by Western Blot analysis. β-Actin was used as a loading control. The experiment was performed on all four hNPC lines and representative images for HUES13-hNPC line are shown. **(C)** Effect of CHIR and XAV (3 μM, 2-3 days) on the mRNA expression of neural genes. qRT-PCR detection of GFAP and MAP2 expression (control – gray bars, CHIR – white bars, XAV – black bars. Values are mean ± SEM. Repeated in 4 hNPC lines, *n*=3/line, *p<0.05, **p<0.01, ANOVA.

First, we assessed the effect of both molecules on the levels of β-Catenin protein in hNPCs (Figure 1B). GSK-3β phosphorylates two different serine residues on the N-terminus of β-Catenin, tagging it for ubiquitination and degradation. TANK indirectly increases β-Catenin levels by de-activating AXIN, an important member of the ‘death complex’ that degrades β-Catenin. Our results confirm that inhibition of GSK-3β by CHIR led to a significant increase in the protein levels of β-Catenin. Conversely, inhibition of TANK by XAV, led to a significant decrease in β-Catenin.

Next, we analyzed whether this short-termed exposure to CHIR and XAV could change the mRNA levels of neural genes reflecting the neurodevelopmental status of hNPCs, including GFAP, MAP2, TUJ1 and TAU (Figure 1C, Figure S1). During formation of mature post-mitotitc neurons and glia, GFAP is reduced in neurons and increased in glial cells, whether MAP2, TUJ1 and TAU are all increased in neurons, as part of the dendritic and axonal cytoskeleton [26]. However, these four genes are all expressed at the hNPC stage, where GFAP corresponds to early hNPC phenotype and MAP2 to advanced hNPC phenotype [8, 24]. Our results show that CHIR and XAV affected GFAP and MAP2 transcriptional levels specifically (Figure 1C), but did not alter TUJ1 and TAU expression (Figure S1). Our results show that short-term exposure to CHIR reduced GFAP in hNPCs (11.4±3.0% of control, p<0.01, *n*=3/line, 4 hNPC lines), but it did not significantly alter the expression of MAP2. On the other hand, XAV had opposite effects in GFAP and MAP2 expression, significantly increasing GFAP (142.0±15% of control, p<0.05, *n*=3/line, 4 hNPC lines) and significantly reducing MAP2 (12.3±9.4% of control, p<0.01, *n*=3/line, 4 hNPC lines). However, since GFAP in this context is an early neural marker, we interpret the XAV-mediated increase in GFAP as representative of reduced neurodevelopment and not enhanced glial differentiation. Taken together, these results suggest that CHIR-mediated inhibition of GSK-3β promotes neural differentiation, whether inhibition of TANK by XAV prevents it.

### Effect of Wnt modulators on morphology, proliferation and survival of hNPCs

To test whether CHIR induced neural differentiation and XAV inhibits it, we subjected the hNPCs to a 7 days-long treatment with CHIR or XAV, and performed a morphological assay which measures neural differentiation (Figure 2A). This morphological assay is based on the premise that, as neural differentiation progresses, cells become bigger and produce more numerous and longer projections, showing an increase in cellular area and perimeter. Indeed, we observed that exposing cells to CHIR for 7 days conferred them a more neuronal phenotype, increasing the number of projections from each cell, and especially their length (Figure 2B). In contrast, XAV elicited the opposite effect, conferring cells with a more hESC-like primitive morphology, lacking projections and demonstrating small round somata. Specifically, CHIR increased the total cell area from 509.1±13.9 μm^2^ to 564.1±15.1 μm^2^ (p<0.05, >50 cells/experiment, *n*=3/line, 4 hNPC lines), while XAV reduced it to 282.2±9.4 μm^2^ (p<0.01, *n* same as above). Their effect on total cell perimeter was similar: CHIR increased cellular perimeter from 106.2±2.0 μm to 120.6±1.7 μm (p<0.05) and XAV reduced it to 71.8±1.5 μm (p<0.01).

**Figure 2.**
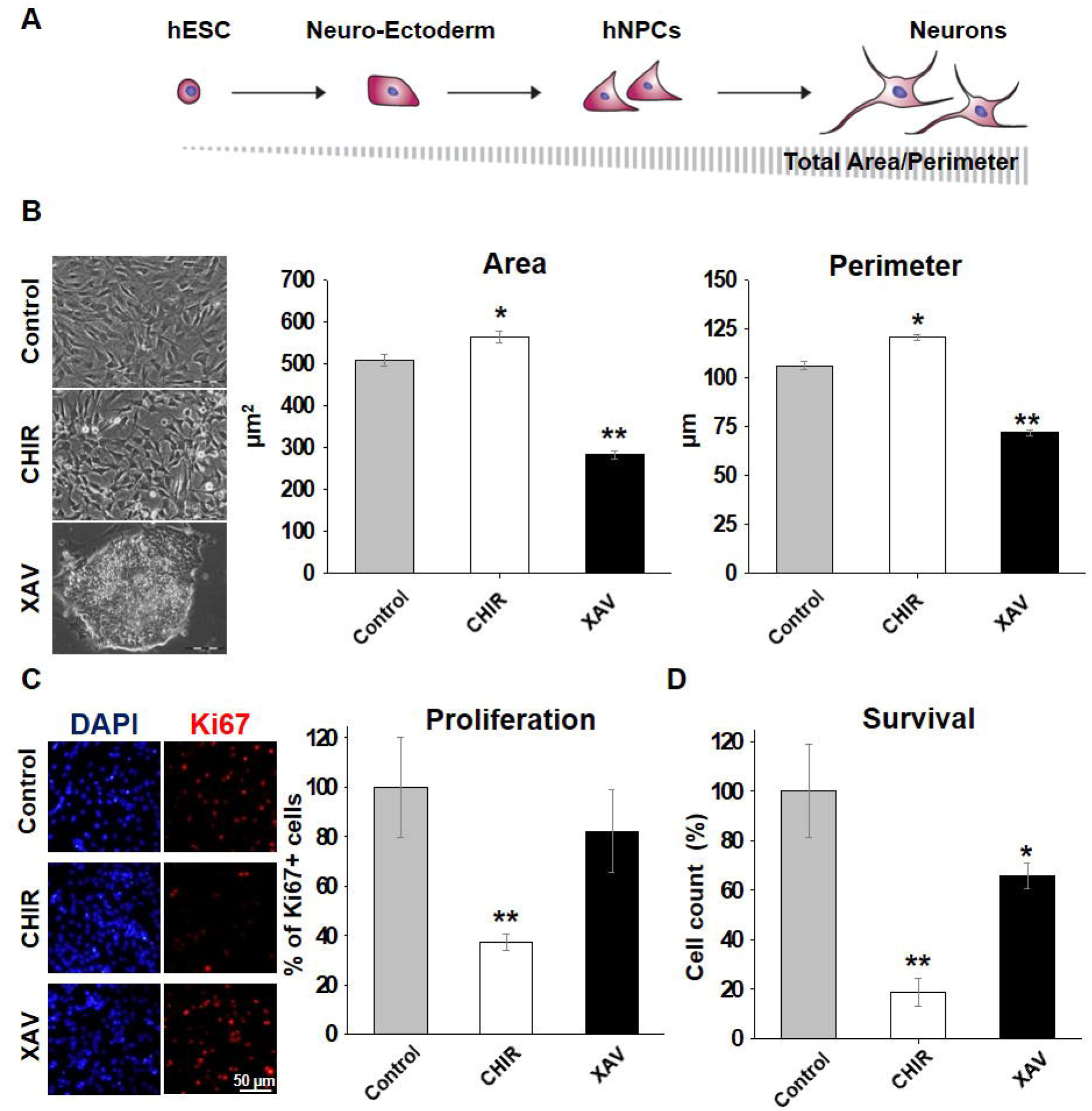
The effect of Wnt modulators on neural fate. **(A)** Schematic presentation of the morphological changes undergoing hESCs during in-vitro neural differentiation, and how they are reflected in the relative change in cell area and perimeter. **(B)** Morphological changes observed in hNPCs following treatment with CHIR or XAV (3 μM, 7 days). The analysis was repeated in all 4 hNPC lines, representative images are shown for of HUES64-hNPCs (left panel). Morphological analysis (right panel) of total area (μm^2^) and total perimeter (μm) carried out for the effect of CHIR (white bars) or XAV (black bars) as compared to control (grey bars). Values are mean ± SEM. Repeated in 4 hNPC lines, n=3/line, *p<0.05, **p<0.01, ANOVA. (**C-D**) Proliferation and survival of hNPCs following exposure to CHIR and XAV (3 μM, 7 days). Cells were stained for DAPI (nuclear staining, blue) and the proliferation marker Ki67 (red). The experiment was repeated in all 4 hNPC lines, representative images shown for HUES6-derived hNPCs. Proliferation was measured as the number of Ki67-positive cells relative to DAPI staining. Survival rate was established by manual count of cells using a standard hemocytometer. Proliferation and survival rate are shown as % of control (control – gray, CHIR – white, XAV – black). Values are mean ± SEM. Repeated in 4 hNPC lines, *n*=3/line, *p<0.05, **p<0.01, ANOVA.

GSK-3β and TANK have a profound effect on regulation of cell proliferation, by regulation of the Wnt/β-Catenin pathway, but also in a Wnt-independent manner. Furthermore, it has been well established that differentiation reduces proliferation, and indeed terminally-differentiated cells, such as adult neurons, are post-mitotic. Correspondingly, undifferentiated or de-differentiated cells, like hESCs or cancer cells, have high rates of proliferation. Therefore, we examined how a 7 days-long exposure to CHIR or XAV could affect proliferation and survival of hNPCs. Quantitative analysis of the expression of the proliferation marker Ki67 showed that treatment with CHIR resulted in a significantly reduced proliferation rate, whereas treatment with XAV did not (Figure 2C). Indeed, the expression of Ki67 was 37.3±3.1% of that found in control hNPC cultures after 7 days of continuous treatment with CHIR (p<0.01, >50 cells/experiment, *n*=3/line, 4 hNPC lines), but it did not significantly differ from the levels of Ki67 expression in hNPC cultures subjected to treatment with XAV. As a means to further confirm this, we performed a survival assay in which cells were seeded at very low densities and treated with CHIR or XAV at the same conditions mentioned before (Figure 2D). This assay confirmed that CHIR evokes a robust reduction in cell survival (p<0.01). This assay also showed that XAV had a statistically significant effect as well (p<0.05), however it was less pronounced than the effect of CHIR. Taken together, these data suggest that CHIR acts as a neuralizing agent in hNPCs, increasing neural differentiation and reducing proliferation. In contrast, XAV acts as a de-differentiation agent, without significantly altering the cell proliferation rate.

### Effect of Wnt modulators on prolonged neuronal differentiation of hNPCs

In order to assess how CHIR and XAV may affect neurogenesis, we induced neuronal maturation in hNPCs and tracked their development for a total of 30 days. The relative increase or decrease in MAP2 and GFAP, is a quantitative tool to directly measure the efficacy of neurogenesis in-vitro, as we and others have previously shown [8, 24]. Therefore, we analyzed the relative protein levels of MAP2 and GFAP using immunostaining assays (Figure 3A,B).

**Figure 3.**
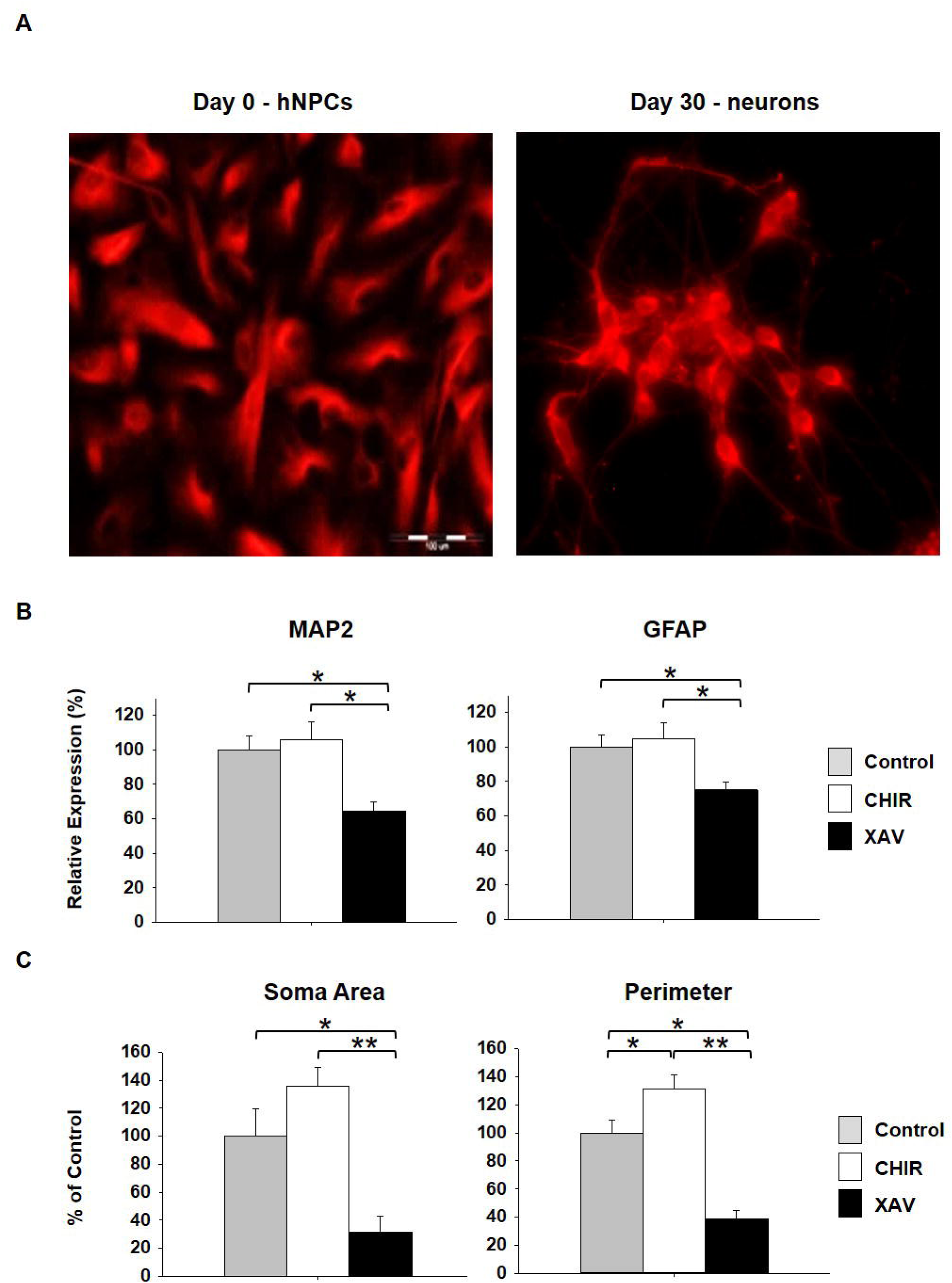
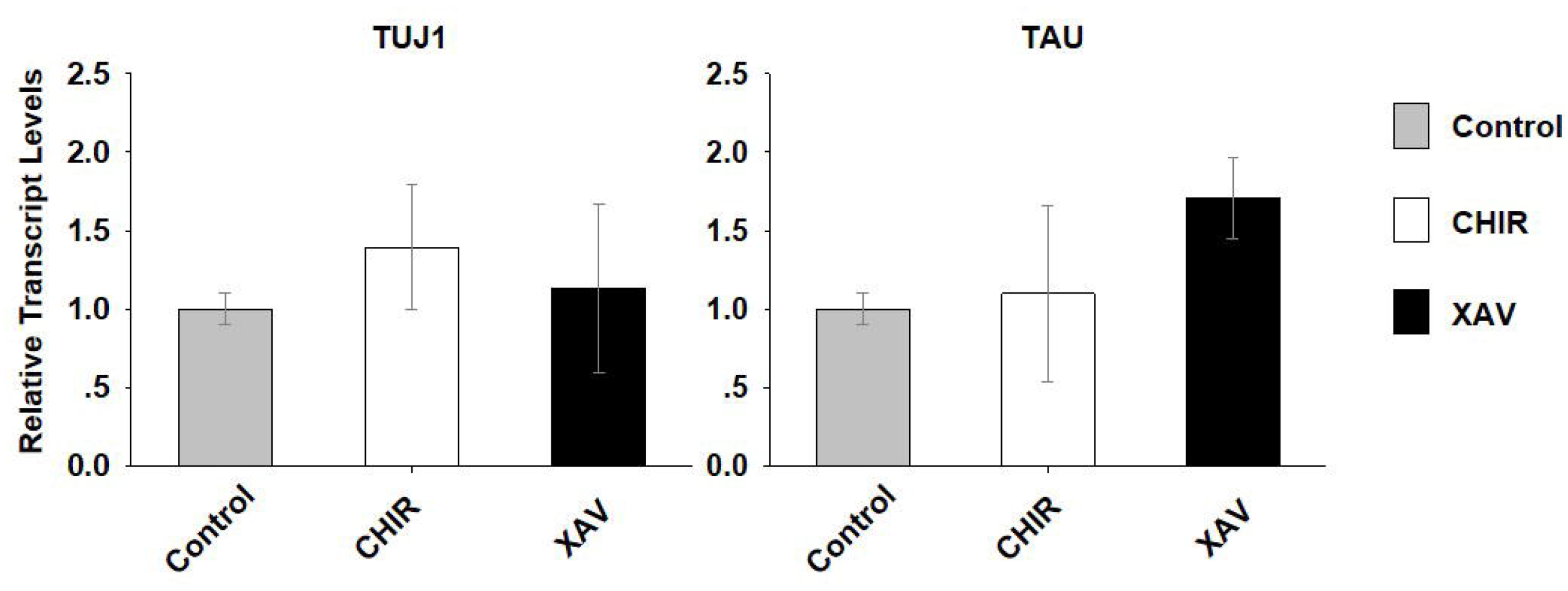
Effect of Wnt modulators on neuronal differentiation of hNPCs. **(A)** Representative images of hNPCs immunostained for MAP2 (red) before beginning of neuronal differentiation (“Day 0”) and following 30 days of neuronal differentiation (“Day 30”). Repeated in all 4 hNPC lines, representative images shown for HUES16 hNPC line. **(B-C)** Effect of CHIR and XAV (3 μM, 30 days) on neuronal differentiation of hNPCs. **(B)** Quantification of MAP2 and GFAP expression relative to DAPI at day 30 of neuronal differentiation of hNPCs (control – gray, CHIR – white, XAV – black). Values are mean ± SEM. Repeated in 4 hNPC lines, *n*=3/line, *p<0.05, ANOVA. **(C)** Relative change in area of cell soma and total cell perimeter at day 30 of neuronal differentiation of hNPCs (control – gray, CHIR – white, XAV – black). Values are mean ± SEM. Repeated in 4 hNPC lines, **n**=3/line, *p<0.05, **p<0.01, ANOVA.

Our results show that by day 30 CHIR did not significantly increase the levels of MAP or GFAP. However, XAV significantly reduced the expression of both markers (MAP2 64.5±5.6% of control, GFAP 75.2±4.6, p<0.05, >50 cells/experiment, *n*=3/line, 4 hNPC lines). Since the effect of CHIR was not reflected in these assays, we analyzed morphological alterations induced by both molecules, as previously described (Figure 2A,B). Since the main morphological hallmark of neuronal differentiation is the development of long and thin neurites, we measured the total area only for somata, whereas the perimeter was measured for the whole cell, including neurites (Figure 3C). Our results show that, after 30 days of treatment, CHIR did not significantly affect the size of cell bodies (135.6±13.7% of control, p>0.05, >50 cells/experiment, *n*=3/line, 4 hNPC lines), but significantly increased their perimeter, reflecting the increase in projection number and length (131.5±9.8% of control, p<0.05). Conversely, XAV robustly reduced both the area of the somas and the total perimeter of cells (area: 31.7±11.4% of control; perimeter: 38.8±6.2% of control, p<0.01). These results confirm the neuralizing effect of CHIR and the de-differentiation effect of XAV during the time-course of neuronal differentiation. All of these measurements were carried-out also at days 10 and 20 of neuronal differentiation, with no significant effects found in neither of those two time-points (data not shown). This suggests that during the initiation of neuronal maturation the activity of GSK-3β and TANK is probably redundant with other molecular mechanisms, and only at later stages they become pivotal in the process.

## DISCUSSION

From the results we have shown in this study we conclude that GSK-3β inhibition by CHIR promotes neural differentiation in hNPCs. Several studies have shown that CHIR induces differentiation into the neural lineage in hESCs, as well as in human induced pluripotent stem cells (hiPSCs) [31–35]. The same was true in primate-derived ESCs and iPSCs [36]. However, in rodent-based models, this was inconsistent. Indeed, two studies reported that in rat-derived ESCs and in mouse embryonic fibroblast, CHIR promoted neural differentiation similar to the human model [37, 38], but two other studies showed that in mouse ESCs CHIR inhibits differentiation, enhancing pluripotency-maintenance mechanisms [39, 40]. This discrepancy between human and murine in-vitro platforms, highlights the importance of the use of human-based models in neurodevelopmental research. Also, it emphasizes the pleiotropic activity of GSK-3β, which can play many different functions that are species and tissue-specific, and depend on developmental status.

In this report we have also shown that XAV induces de-differentiation of hNPCs towards a more primitive developmental stage. However, to the best of our knowledge, no other similar reports have been produced so far, in which the effect of XAV was analyzed in hNPCs derived from hESCs or hiPSCs. In a human neurobolastoma cell line XAV was found to induce apoptosis by blocking Wnt signaling [41]. In zebrafish embryos, XAV inhibited retinal development [42]. Using mouse Epiblast-derived stem cells, XAV increased the development of forebrain precursors in one report [43], but inhibited differentiation and enhanced pluripotency in another report [44]. Finally, in mESCs, it was shown that XAV promotes differentiation into cardiomyocytes [45]. These discrepancies could be attributed to the model used in each study, and to whether in those tissues TANK activity affects the regulation of the Wnt signal only, or not is involved in other currently unknown mechanisms. Here too, similar to GSK-3β, the pleiotropy of TANK activity could lead to mixed interpretations of the results

Our data suggest that the neuralizing effect of CHIR and the anti-neuralizing effect of XAV on hNPCs, is probably due to their involvement in the Wnt signaling pathway and their opposite effect on β-Catenin levels. This is because CHIR and XAV had opposite effects on cell proliferation as well as on neural differentiation. However, more research is needed to determine which specific functions of GSK-3β and TANK are Wnt-dependent and which are Wnt-independent in the regulation of hNPCs developmental fate.

Finally, we have shown here that hNPCs derived from hESCs are a valuable tool for in-vitro human-based drug screening studies. As briefly reviewed here, there are striking discrepancies between human-based in-vitro models and animal-based models. Finding the molecular basis for these differences is crucial to improve basic knowledge as well as to improve future therapeutic strategies.

