## Supplemental Tables and Figure. for "Neural differentiation is increased by GSK-3β inhibition and decreased by tankyrase inhibition in human neural precursor cells"

**Supplementary Table 1**

| Gene | Reverse | Forward |
|---|---|---|
| *GAPDH* | atacgaccaaatccgttgactc | agccacatcgctcagacacc |
| MAP2 | cattggcgcttcggacaag | ctcagcaccgctaacagagg |
| GFAP | aggtccatgtggagcttgac | gccattgcctcatactgcgt |
| TUJ1 | tttttgctcgcctcaaggtatgt | gggcgcattccaacctt |
| TAU | tgccatgttgagcaggacta | tcacttttacagcaacagtcagtg |

**<u>Table S1</u>. Primers (oligos) used for quantitative Real Time PCR.**

Sequences of all primers (oligo-DNAs) used for quantitative Real Time PCR assays (Sigma).

**Supplementary Table 2**

| Target | Host | Provider and Cat. # |
|---|---|---|
| GFAP | Mouse | Millipore - MAB360 |
| MAP2 | Rabbit | Santa Cruz - sc-20172 |
| β-Catenin | Rabbit | Santa Cruz - sc7199 |
| β-Actin | Mouse | Abcam – ab8224 |

**<u>Table S2</u>. Primary antibodies used for immunostaining and western blot**

Full list of primary antibodies used in immunostaining assays and western blot analysis. For immunostaining, primary antibodies were detected using Cy2-conjugated sheep anti-mouse or Cy3-conjugated goat anti-rabbit secondary antibodies (Jackson Labs). For western blot, primary antibodies were detected using HRP-conjugated goat anti-mouse or HRP-conjugated goat anti-rabbit secondary antibodies (Jackson Labs). Working concentrations of primary and secondary antibodies were as recommended by manufacturer.

**Supplementary Figure 1.**

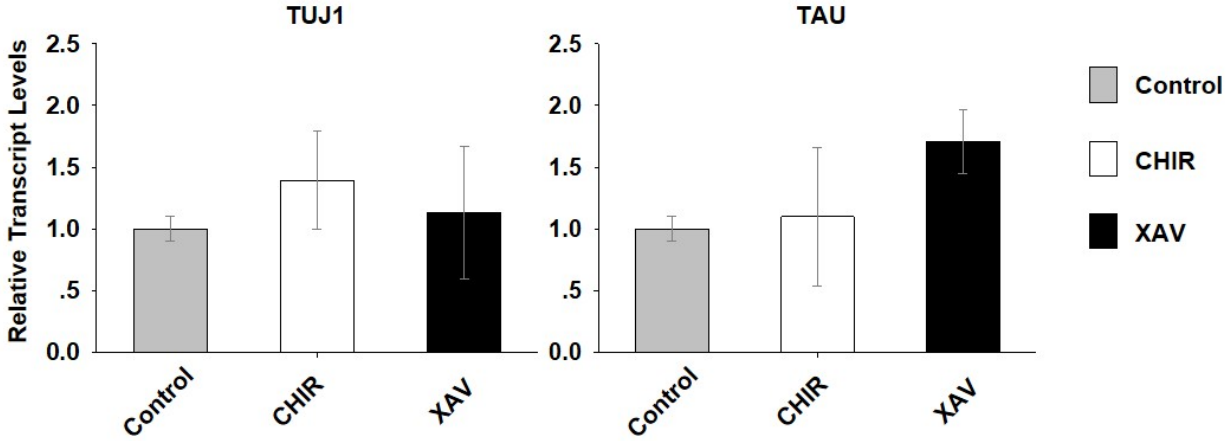


**Figure S1. qRT-PCR analysis of TUJ1 and TAU expression.**

Effect of CHIR and XAV (3 µM, 2-3 days) on the mRNA expression of neural genes. qRT-PCR detection of TUJ1 (left) and TAU (right) expression (control – gray bars, CHIR – white bars, XAV – black bars. Values are mean ± SEM. Repeated in 4 hNPC lines, *n*=3/line, *$p<0.05$, **$p<0.01$, ANOVA, all differences were non-significant ($p>0.05$).
